# The evolutionary advantage of toxin production among cyanobacteria, the oldest known organisms on Earth

**DOI:** 10.1101/615542

**Authors:** Ravi José Tristão Ramos, Crina-Maria Ionescu, Jaroslav Koča

## Abstract

Cyanobacteria produce toxic secondary metabolites for reasons hitherto unclear. Using a phylogenetic approach that accounts for the high complexity of biosynthetic gene clusters (full or partial inversions, variable length, different number of genes, non-orthologues), we analyzed the sequences of 76 biosynthetic gene clusters covering 19 cyanotoxins. The phylogenetic tree of biosynthetic gene clusters branches first according to the bioactivity of the toxic metabolite (molecular target in another organism), then according to the chemical class and chemical structure of the secondary metabolite, and finally according to the organism and area of origin. The bioactivity of a toxic metabolite can be deduced directly from the nucleotide sequence of the biosynthetic gene cluster, without needing to examine the enzymes themselves or to measure expression levels. Bioactivity may have been the primary driving force behind the diversity of secondary metabolism in cyanobacteria. This genetic machinery evolved to facilitate three specific survival strategies acting separately or in tandem, with dominant cyanobacteria possessing the genetic machinery to support all three strategies. Transmembrane (direct) toxicity targeting ion channels, intracellular (indirect) toxicity targeting cell-cycle regulation, and digestion inhibition targeting proteases may have provided the survival advantage underpinning the evolutionary success of both cyanobacteria and their early symbiotic hosts.

## Introduction

Bioactive secondary metabolites, which are products of pathways not directly involved in growth, development, or reproduction, often serve complex functions that aid survival and increase dominance of an organism in a given ecological niche. Snake venoms are probably the most famous examples of bioactive secondary metabolites, though not nearly the most versatile. Bacteria, fungi, and plants all exhibit an impressive array of secondary metabolites, with cyanobacteria being the earliest organisms to develop such capabilities. Cyanobacteria are unicellular photosynthetic prokaryotes populating marine, fresh-water, and land ecosystems, which drove the rapid oxygenation of Earth’s atmosphere and are recognized as the ancestors of plastids^1^. Cyanobacteria continue to exert a crucial environmental impact because they produce about 25% of all carbohydrates on Earth and perform massive nitrogen fixation together with sequestering trace metals and phosphorous from the environment, thus fueling most contemporary food chains. In addition, many cyanobacteria produce unique secondary metabolites (typically, peptides and alkaloids) via non-ribosomal peptide synthesis (NRPS) and/or polyketide synthesis (PKS), with complex environmental, economic, and health impact^2^.

The exact function of cyanobacterial secondary metabolites remains largely unknown^2^ despite the fact that a substantial portion of the genome can be dedicated to this secondary metabolism^3^. These bioactive compounds are often referred to generically as cyanotoxins because toxic metabolites are found in great quantities during cyanobacterial blooms that kill local aquatic fauna. However, not all secondary metabolites of cyanobacterial NRPS/PKS are especially toxic, and some even have strong potential for use as pharmaceuticals, pesticides, or biofuels^4,5^. It is strongly suspected that production of toxins and other bioactive peptides is linked to the evolution of certain cyanobacterial species or niche populations^2,4,6^. To date, the following potential roles have been suggested for bioactive NRPS/PKS products: (i) photoprotection due to absorption of harmful ultra-violet radiation; (ii) facilitating the acquisition or more efficient use of available but limiting resources; (iii) impeding growth or resource use of competitors such as green algae or other cyanobacteria; and (iv) limiting predation by inducing toxicity in organisms predating on cyanobacteria^2,4,6^. While the role of cyanobacterial metabolites in photoprotection, resource monopolization, and habitat dominance has been demonstrated experimentally, hypotheses regarding the anti-predation role are not entirely consistent^2,4,6^. Specifically, some studies have provided evidence of co-evolutionary adaptation between toxin-producing cyanobacteria and its natural predator, zooplankton^7^, while others have suggested that the oldest cyanotoxin genes appeared before zooplankton^8^. Thus, to clarify the evolutionary role of bioactive secondary metabolites in cyanobacteria, it is necessary to take a second look at the genetic machinery responsible for the production of such compounds.

It is currently possible to study secondary metabolism in great detail using dedicated high-throughput omics technologies that enable to quantify the levels of entire pools of transcripts, proteins, and metabolites in a given cell, in a particular tissue, and even in environments populated by multiple species. However, substantial information regarding the nature of the final metabolite can be gained simply from examining the genes involved^9^. Moreover, it is currently possible to predict the chemical class of a metabolite using only the translated sequence of the genes coding for the set of enzymes responsible for metabolite production, together with information from dedicated databases covering known genes, enzymes, pathways, and products of secondary metabolism^10^. Going one step further, one may wonder whether the nucleotide sequences biosynthetic gene clusters themselves already contain the key information regarding the impact of the secondary metabolite in the environment (i.e., bioactivity). Thus, we set out to examine the relationship between nucleotide sequence similarity and bioactivity of secondary metabolites produced by complex gene clusters performing NRPS/PKS in various species of cyanobacteria. Understanding these relationships may provide insight into the potential contribution of toxin-producing gene clusters in the overwhelming evolutionary success of cyanobacteria and in their role as promoters of life diversification on Earth.

However, analyzing cyanobacterial genomes is challenging because cyanobacteria, which are the earliest known organisms on our planet, have accumulated a huge amount of lateral gene transfer and gene shuffling events over geological time^11,12^. In fact, the phylogenetic relationships among cyanobacterial species remain an important topic of debate^13–15^. Regarding the genetic basis of cyanotoxin production, the challenge lies in the high variability of the biosynthetic gene clusters. Tens of secondary metabolites with different structure and bioactivity may be produced via NRPS/PKS within the same genus, yet different strains of the same species might not produce the same metabolites^6,12,13^. Corresponding NRPS/PKS gene clusters in different species or even in different strains of the same species differ regarding the presence, order, direction, and length of genes, which may or may not be orthologous^16,17^. It remains unclear just how much horizontal gene transfer occurs from different strains and different species of cyanobacteria, as well as from non-cyanobacterial species. For example, upon phylogenetic analysis of cylindrospermopsin-producing genes, *Cylindrospermopsis raciborskii* from Spain grouped together with Tunisian and American strains and not with other European strains^18^ even though *C. raciborskii* is reported to form a tight monophyletic group with specific continental distribution in terms of normal phylogenetic markers such as the 16s rRNA gene^19,20^. In the case of the saxitoxin-producing gene cluster, mainly vertical transfer congruent with 16s rRNA evolution was suggested for some components (e.g., genes encoding for aminotransferase or histidine kinase), whereas horizontal gene transfer has been suggested for other components (e.g., genes encoding for polyketide synthase or amidinotransferase)^21,22^. A recent comparative genomics analysis highlighted evidence for massive gene shuffling within biosynthetic gene clusters across the *Cyanobacteria* phylum, with potential crossover to and from non-cyanobacterial species, and without coherence with species phylogeny^12^. It is clear that a multi-gene phylogenetic approach is required here, but such analyses involving highly divergent non-orthologues have been very challenging^12,23^. Therefore, it is desirable to apply a unified measure of sequence similarity that can be used to compare complex stretches of DNA along entire biosynthetic gene clusters even in the absence of further information regarding evolutionary mechanisms.

To tackle this complex problem, we employed a phylogenetic approach that bridges alignment-based and alignment-free phylogenomics, facilitating the analysis of long stretches of DNA containing orthologous and non-orthologous sequences with high or uneven divergence, gene shuffling, and inversions. We found that the sequence of complex biosynthetic gene clusters involved in the secondary metabolism of cyanobacteria contains key information regarding the bioactivity of the final product. These relationships were apparent directly from the nucleotide sequence of the biosynthetic gene clusters, without the need to examine the enzymes encoded by the genes in each cluster or to measure protein or RNA expression levels, suggesting that bioactivity may have been the primary driving force behind the diversity of the genetic machinery for secondary metabolism in cyanobacteria. Finally, we explored the implications of our findings in the context of cyanobacterial evolution. Our findings suggest that the genetic machinery responsible for secondary metabolism in cyanobacteria may have evolved to facilitate specific survival strategies in addition to a potential role in resource management. Transmembrane toxicity targeting ion channels, intracellular toxicity targeting cell-cycle regulation, and digestion inhibition targeting proteases may have provided the survival advantage underpinning the evolutionary success of both cyanobacteria and their early symbiotic hosts.

## Results

### Key information regarding the bioactivity of a secondary metabolite can be found directly within the nucleotide sequence of its biosynthetic gene cluster

We surveyed the NCBI Nucleotide database and obtained the sequences of 76 gene clusters responsible for NRPS/PKS in cyanobacteria (approximately 3000–64000 bp each), covering 19 different products (cyanotoxins). The complexity of these gene clusters is illustrated in Supplementary Table S1, which summarizes the organism of origin, the number and order of genes in each cluster, and the nature of the secondary metabolite produced by the enzymes encoded by the gene cluster.

To account for the complex nature of these NRPS/PKS gene clusters, we employed a phylogenetic approach built around local pairwise sequence alignment (LPSA), which is commonly used to identify, categorize, annotate, and filter out gene or protein sequences obtained from one or more species. LPSA has been shown to perform well for phylogenetic analyses involving multiple genes and even whole genomes of viruses, eukaryotes, and bacteria, including cyanobacteria^24–28^. The LPSA-based approach used in this study (https://crocodile.ncbr.muni.cz/) involves three steps, namely: (i) break each sequence into subsequences; (ii) using the Basic Local Alignment Search Tool (BLAST)^29,30^, perform local alignment of each pair of subsequences generated for the entire dataset; and (iii) using the BLAST report for subsequences (percentage of identities on matches, subsequence and alignment lengths), build a matrix that reflects the distance between each pair of sequences in the original dataset (full methodological details are provided in the Methods). The information contained in the distance matrix is used to generate a phylogenetic tree in which branch lengths are derived from the average pairwise similarity of subsequence alignments.

Upon analyzing the biosynthetic gene clusters using this LPSA-based approach, we obtained a phylogenetic tree describing the sequence similarity relationships among the 76 gene clusters (Figure 1). The phylogenetic tree of biosynthetic gene clusters branches first according to the bioactivity of the secondary metabolite, which is given by the molecular target of the cyanobacterial metabolite in another organism. Specifically, gene clusters producing ion channel modulators group separately from those producing phosphatase inhibitors and from those producing protease inhibitors. Furthermore, gene clusters producing neurotoxins targeting voltage-gated ion channels (saxitoxins) group separately from gene clusters producing neurotoxins targeting ligand-gated ion channels (anatoxins). These results confirm that the biosynthetic genes themselves already contain key information regarding the impact of the secondary metabolites in the environment (i.e., bioactivity). Furthermore, these findings suggest that the bioactivity of the secondary metabolites may be the primary driving force behind the diversity of the genetic machinery for secondary metabolism in cyanobacteria.

**Figure 1.**
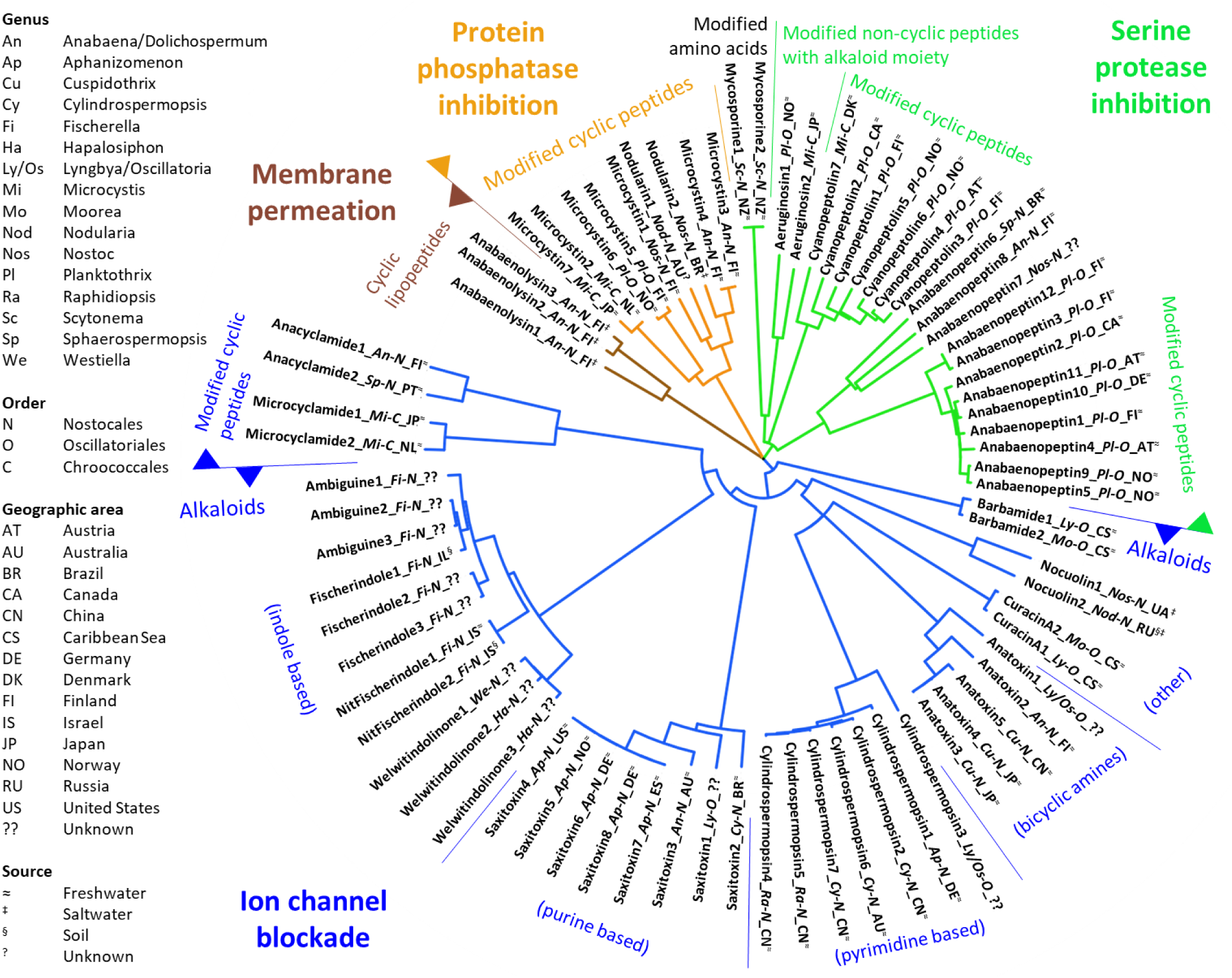
Relationships between nucleotide sequence similarity of biosynthetic gene clusters and characteristics of secondary metabolites produced. The phylogenetic tree is organized first according bioactivity (molecular target in other organisms), then according to chemical structure, and finally according to organism of origin and geographical area of origin (full details in Supplementary Table S1). Branch lengths do not reflect evolutionary time but are derived from the average pairwise similarity of subsequence alignments. For visualization purposes, the clades are colored according to the confirmed molecular targets of metabolites making up relevant nodes. Full methodological details are provided in the Methods.

### Bioactivity, rather than similarity of chemical structure, is the main explanation behind genetic diversity in the secondary metabolism of cyanobacteria

The next levels of branching of the phylogenetic tree correspond to the chemical class and chemical structure of the secondary metabolite, which are dictated by the biosynthetic enzymes and the order of biosynthetic reactions. Specifically, gene clusters producing cyclic peptides generally group separately from those producing alkaloids. Furthermore, among gene clusters producing indole-based alkaloids, there is distinct grouping according to whether the bioactive metabolite contains a nitrile group or an isonitrile group (NitFischerindole vs. Fischerindole in Figure 1). However, gene clusters producing anacyclamide and microcyclamide group with clusters producing indoles and not with clusters producing other modified cyclic peptides. Furthermore, it should be noted that most secondary metabolites produced via NRPS/PKS are modular compounds, and it is currently recognized that variability in one or more substituents leads to different target specificity. For example, the affinity for genetic variants of the same ion channel differs across saxitoxin analogs^31^, whereas the specificity for serine proteases with trypsin- and thrombin-like activity differs among aeruginosins^32^. Thus, the grouping of biosynthetic gene clusters according to the chemical structure of the secondary metabolite likely reflects the bioactivity of the metabolite rather than its specific chemical structure.

Indeed, gene clusters producing nodularins grouped closely together with those producing microcystins. While both nodularins and microcystins are cyclic peptides, nodularins typically contain five amino acid residues, whereas microcystins contain seven; however, the biological activity of nodularins and microcystins is equivalent, as these metabolites have the same molecular targets, the same binding site on their molecular target, and almost the same interactions involved in binding with the molecular target^16,33^. Furthermore, despite the fact that aeruginosins are linear and contain an indole part in addition to the oligopeptide chain, gene clusters producing aeruginosin grouped together with those producing cyclic peptides that share the same molecular target in other organisms (i.e., serine proteases)^32^. These findings suggest that genetic diversity in the secondary metabolism of cyanobacteria is explained mainly in terms of bioactivity rather than in terms of chemical structure similarity.

### Non-uniform horizontal gene transfer of biosynthetic gene clusters of the secondary metabolism

The next level of branching of the phylogenetic tree corresponds to the genus/species and geographical area of origin. For example, saxitoxin-producing gene clusters originating from *Aphanizomenon* species group separately from clusters originating from other genera; similar findings are noted for anabaenopeptin-producing clusters originating from *Planthotrix* species, as well as for cyanopeptolin-producing clusters originating from *Planthotrix* species. Other relationships are less well defined. Cylindrospermopsin-producing cluster 7, which comes from a Chinese *C. raciborskii* strain, groups closer to clusters 4 and 5, which come from Chinese *Raphidiopsis curvata* strains, than with cluster 2, which comes from a different Chinese *C. raciborskii* strain and groups closer to cluster 6, which comes from an Australian *C. raciborskii* strain. The anabaenopeptin-producing cluster 8, which originates from a Finnish *Anabaena* species, groups separately from clusters 1, 3, and 12, which come from Finnish *Plankthotrix* species; on the other hand, cluster 3 groups close to cluster 12, as they both come from the same Finnish *Plankthotrix aghardii* strain, while cluster 1 from another Finnish *P. aghardii* strain groups closer to clusters from non-Finish *Plankthotrix* species. These findings confirm that some biosynthetic gene clusters may transfer more readily than others across species or strains, likely aided by geographical proximity or contamination events, while other signs of recombination and gene duplication are likely related to vertical inheritance^12^.

## Discussion

### Survival strategies based on secondary metabolism

Based on the organization of the phylogenetic tree of biosynthetic gene clusters, three main strategies with potential survival benefit can be distinguished, namely transmembrane (direct) toxicity due to ion channel blockade, intracellular (indirect) toxicity due to protein phosphatase blockade, and digestion inhibition due to serine protease blockade (Figure 2). Various branches of secondary metabolism contributing to each strategy likely provide back-up or some form of specialization to facilitate the strategy. Direct and indirect toxicity may act separately or in tandem, with the effects of indirect toxicity taking longer to manifest, requiring facilitation by a membrane permeabilization agent, and being mainly aimed at organisms of the animal kingdom. Digestion inhibition would serve as a bail-out strategy in case the cell is ingested, not particularly aiming to kill the culprit organism; in other words, serine protease inhibition by cyanobacterial toxins likely arose as a purely protective mechanism, which is fundamentally different from the role of serine protease inhibitors produced by animals (e.g., ticks)^34^. Dominant cyanobacteria such as *Planktothrix*, *Anabaena*, and *Microcystis* have the genetic machinery to support all three strategies.

**Figure 2.**
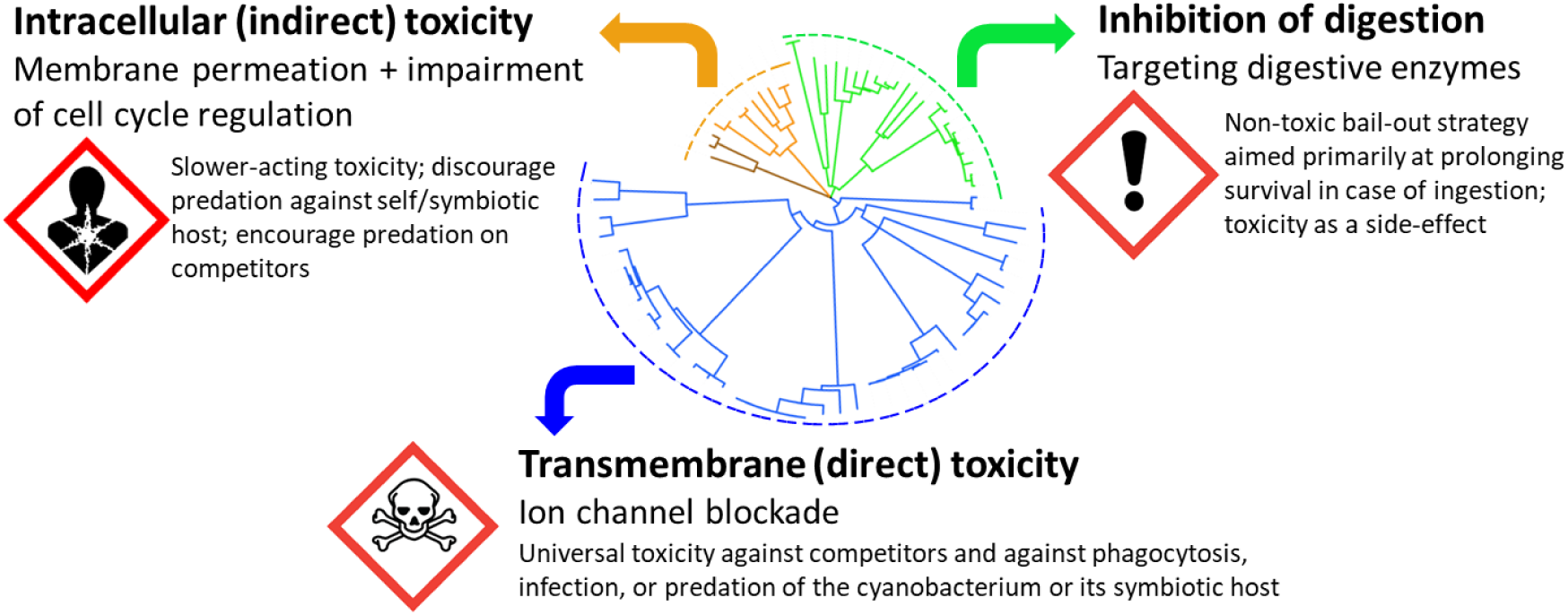
Evolutionary advantage of toxin production by cyanobacteria. Three survival strategies can be distinguished. Dominant cyanobacteria such as Planktothrix, Anabaena, and Microcystis have the genetic machinery to support all three strategies.

***Direct toxicity*** can be achieved by modulating the activity of voltage- and ligand-gated ion channels, which are ubiquitous across all life domains. Such mechanisms, which often involve alkaloid metabolites with relatively broad specificity but differential affinity, can be used effectively against competitors and against phagocytosis, infection, or predation by other organisms^35–37^. Our findings support the role of the neurotoxin saxitoxin as a modulator of voltage-gated ion channels^2^ and further suggest that indole-based alkaloids and perhaps cyclamides may share such targets. The indoles welwitindolinone and fischerindole have rarely been studied. Nevertheless, related indole-based alkaloids produced by the genus *Fischerella* are suspected to act as ion channel modulators similar to saxitoxin^38^. Furthermore, the indole moiety itself is a known modulator of voltage-dependent gating of potassium channels^39^. Cyclamides are cyclic oligopeptides produced via the ribosomal pathway and heavily functionalized via post-translational modifications^40^. While anacyclamide and microcyclamide have not been studied for their interaction with ion channels, the thiazoline moiety commonly encountered in such cyclamides has been reported to be critical for the toxicity of kalkitoxin^41^, another cyanotoxin that targets voltage-gated ion channels^42^. At the same time, our findings support the role of the neurotoxin anatoxin as a modulator of ligand-gated ion channels^2^, and further suggest that cylindrospermopsin, curacyn, nocuolin, and perhaps barbamide may share such targets. While cylindrospermopsin is typically referred to as a hepatotoxin, its neurotoxicity has been confirmed in fish, with the nicotinic acetylcholine receptor as the suspected target (i.e., the same target as that of anatoxin), though binding seems reversible^43^. The interaction of curacin A with ligand-gated ion channels has not been tested to date. Curacin A-like compounds produced by the same species (tropical *Lyngbia majuscula*) were shown to induce neurotoxicity in rat neurons via interaction with ligand-gated ion channels, though curacin A itself did not have such effect in rat neurons^44^. As curacin A is toxic to shrimp^36^, the difference in toxicity (shrimp vs. rat neurons) might be related to differences in specificity towards the target molecule. Finally, nocuolin A and barbamide have not been studied for potential neurotoxicity or interaction with ion channels. Nocuolin A is a natural oxadiazine, a class of compounds shown to block sodium channels and exert neurotoxic effects in cockroaches^45^. Barbamide is a strong molluscicide, though its mechanism of action has yet to be determined. Given that barbamide contains thiazole and trichloromethyl, two pharmacophores known to interact with ion channels^45,46^, it is likely that barbamide also targets ion channels.

***Indirect toxicity*** can be achieved by modulating intracellular signaling. Signaling processes involved in growth and reproduction rely on recognition of specific protein phosphorylation patterns created by the interplay of protein kinases and protein phosphatases. Microcystins/nodularins, which block protein phosphatases, interfere with the cell cycle leading to abnormal mitosis and even cell death. In addition to actively deterring predation^35,47^, microcystins indirectly support dominance of the producing cyanobacterium by promoting preferential predation on competing organisms^48^. It has been suggested that nodularin synthetase genes are derived from microcystin synthetase genes^8^. While the pentapeptide nodularin and the heptapeptide microcystin are similar in structure and bioactivity, neither has a strong effect on its own. Instead, anabaenolysin may potentiate the action of microcystins/nodularins by permeating cholesterol-based membranes^49^. In microcystin/nodularin producers, biosynthetic clusters for anabaenolysin may have evolved as an adaptation to the evolution of animal organisms feeding on the symbiote. A single cyanobacterial strain may produce many anabaenolysin variants^50^, probably as an adaptation to the diversification of marine life.

***Digestion inhibition*** can be achieved by modulating the activity of proteases, which protects ingested cells from being degraded. As such digestive enzymes have highly efficient catalytic sites, inhibition requires high specificity, which is provided by peptide-based metabolites. Aeruginosin, anabaenopeptin, and cyanopeptolin are serine protease inhibitors, typically targeting the digestive enzymes trypsin and/or chymotrypsin, as well as similar proteases involved in the coagulation pathway. The same cyanobacterium may possess the genetic machinery to synthesize more than one type of serine protease inhibitors and there is substantial though not complete overlap in the specificities of aeruginosin, anabaenopeptin, and cyanopeptolin^6,32^, suggesting that these biosynthetic gene clusters may provide mutual back-up for inhibiting digestion in high-level organisms predating/grazing on cyanobacterial symbiotic hosts. In this context, interference with the coagulation pathway may not be evolutionarily motivated in cyanobacteria, but rather a consequence of the similarity between digestive proteins and coagulation pathway proteins. At the same time, the evolution of genetic machineries to produce serine-protease inhibitors in the form of cyanobacterial metabolites may have contributed to the diversification of tryptic enzymes in the animal kingdom.

### Implications for the study of cyanobacterial evolution

While it is agreed that microcystin- and saxitoxin-producing gene clusters are likely the oldest toxin-producing gene clusters in cyanobacteria, it is believed that the ability to produce microcystin and saxitoxin has been lost repeatedly across multiple lineages of cyanobacteria and even among strains of the same species, suggesting that there may have been neither negative nor positive pressure for such biosynthetic genes, or that such pressure was circumstantial to some clades^6,8,22^. However, others have pointed out that the genomes of South American saxitoxin-producing *C. raciborski* strains contain vestiges of cylindrospermopsin-producing genes, whereas no vestiges of saxitoxin-producing genes are seen in the genomes of the cylindrospermopsin-producing strains^51^. Our findings suggest that saxitoxin/cylindrospermopsin and microcystin/nodularin serve parallel survival strategies (transmembrane vs. intracellular toxicity, respectively; Figure 1). Due to its simplicity, accessibility, and fast acting mechanism, transmembrane toxicity by modulation of ion channels likely represents an older mechanism^52^.

### Limitations

The phylogenetic analysis approach applied in the present study is conceptually similar to previous efforts employing average sequence similarity^24–28^, in that it relies on obtaining LPSA similarity scores for certain pieces of the genome, averaging the score for each full sequence analyzed, and creating a distance matrix based on the average similarity scores. However, unlike other LPSA-based phylogenetic approaches^28^, the method used here does not employ information regarding evolutionary timelines. For this reason, the branch lengths of the phylogenetic tree obtained in this study are not directly representative of evolutionary time but of sequence similarity, for which the evolutionary rate may vary throughout geological time and environment. On the other hand, cyanobacteria evolution would have been heavily influenced by their symbiotic relationships, which may preclude the application of current assumptions regarding evolutionary rate^53,54^. Furthermore, we could not infer ancestral sequences because this LPSA-based approach obtains distance matrices based solely on average similarity scores, which inevitably results in partial loss of alignment information. Finally, while many cyanobacterial metabolites have been characterized^2^ and many sequences potentially corresponding to NRPS/PKS enzymes have been detected^55^, the coding sequences currently available for other such metabolites did not fit the inclusion criteria for this study. As more data accumulate, it will become possible to perform an all-encompassing phylogenetic analysis of cyanobacterial biosynthetic gene clusters and shed further light into the potential biological, ecological, and evolutionary role of cyanotoxins.

### Scope for further research

The high level of organization of the phylogenetic tree of biosynthetic gene clusters illustrates the depth of information that can be obtained using LPSA combined with a simple method of overall sequence similarity. It is notable that we were able to obtain such information without having to evaluate non-coding regions, RNA/protein expression levels, or metabolomics profiles. This suggests that, when attempting to understand the relationships among complex phenotypes in prokaryotes, it may not be necessary to build one tree per gene or even one tree per gene cluster. We propose that the “sequence-structure-function” paradigm, typically discussed in the context of proteins, can be expanded from the nucleotide sequence of the gene cluster to the bioactivity of the metabolite synthesized by the enzyme ensemble encoded by the gene cluster.

Thus, our findings can also be interpreted in the context of the “It’s the song not the singer” (ITSNTS) theory, which proposes that the true units of evolutionary selection are not the organisms themselves (i.e., the singers in the ITSNTS theory) but the processes implemented by genes, cells, species, or communities (i.e., the song in the ITSNTS theory)^56^. In this context, we may speak about the bioactivity of the biosynthetic gene cluster (i.e., the song in the ITSNTS theory) as an independent entity of a biosystem, driving the evolution of organisms or communities carrying such biosynthetic genes or interacting with the metabolites synthesized by the genes (i.e., the singers in the ITSNTS theory). Nevertheless, looking only at the gene cluster sequence is not sufficient to clarify whether and how much of a given toxin will be produced under different environmental conditions. While it is generally accepted that recent cyanobacterial invasion, proliferation, and ecological impact are strongly affected by climate change and environmental changes caused by human activities^57,58^, it remains unclear what triggers abundant production of toxic metabolites by otherwise silent gene clusters, though it is quite evident that such triggers are multi-faceted^2^. Further study is warranted to clarify whether or how secondary metabolites could enhance resource utilization or temperature and chemical resistance in cyanobacteria. Alternatively, since the feeding behavior of grazers and predators is sensitive to environmental changes, the fact that toxin production in cyanobacteria is triggered by environmental changes may simply reflect the association between such changes and feeding behavior of predators. Finally, the survival strategies proposed in this study may not have been the original ecological roles of cyanotoxins, as these early roles would have been lost or replaced with new functions over evolutionary time, reflecting the change in local and planet-wide ecological context. It is important to keep in mind that cyanobacteria are often encountered as endosymbionts of corals, diatoms, dinoflagellates, seagrass, and sponges. Thus, survival benefits conferred by bioactive secondary metabolites may apply to the cyanobacterium itself or to its symbiotic host. For example, while there may be negative selective pressure for microcystin-producing gene clusters within the Cyanobacteria phylum^8^, such genes may provide benefit to the symbiotic host^59^. Production of toxic metabolites by cyanobacteria may have promoted the adaptation of some animals to consuming a diet contaminated with toxin-producing cyanobacteria and even carrying the cyanobacteria as a symbiont; for example, all eight sodium channel variants in the pufferfish genome harbor mutations rendering them resistant to saxitoxin^60^. The impact of the survival strategies discussed here should therefore be considered in the context of these symbiotic relationships. Furthermore, future genomic studies on cyanobacteria should take into account the symbiotic host, as gene transfer potential may be substantially altered for cyanobacteria living as symbionts^54^.

### Conclusion

The bioactivity of a toxic metabolite can be deduced directly from the nucleotide sequence of the biosynthetic gene cluster, without needing to examine the enzymes themselves or to measure expression levels. Bioactivity may have been the primary driving force behind the diversity of secondary metabolism in cyanobacteria. This genetic machinery evolved to facilitate three specific survival strategies acting separately or in tandem, with dominant cyanobacteria possessing the genetic machinery to support all three strategies. Transmembrane (direct) toxicity targeting ion channels, intracellular (indirect) toxicity targeting cell-cycle regulation, and digestion inhibition targeting proteases may have provided the survival advantage underpinning the evolutionary success of both cyanobacteria and their early symbiotic hosts.

## Methods

### Description of the LPSA-based approach for phylogenetic analysis

a. Definitions
  ***S***: A sequence to align
  ***L***: The length of a given sequence
  ***s***: A subsequence of ***S***
  ***σ_i_***: The subsequence of ***S_i_*** with the highest similarity to a certain subsequence ***s*** ***l***: Length of a subsequence ***s***
  ***f***: Frame shift from position 1 of a sequence ***S*** ***n***: Number of frames ***f*** within a subsequence ***s***

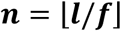
  ***N***: Number of subsequences in ***S***, including the subsequences from different frame shifts ***f***

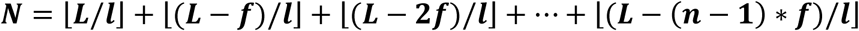
  ***N_0_***: Number of subsequences in ***S_i_*** for which no subsequence in ***S_j_*** was found during the alignment
  ***M***: The distance matrix for all sequences ***S***
b. Description
  The approach involves three steps described schematically below for a dataset of three sequences.
    i. Break each sequence ***S_i_*** (for 3 subsequences: ***i, j*** ∊ {***a, b, c***}) into ***n_i_*** subsequences of a chosen length ***l***, equal for all ***i***
    ii. Perform pairwise local alignment (in this case, using BLAST) of all subsequences ***s_i_*** vs all subsequences ***s_j_***, ***i*** ≠ ***j***
    iii. Calculate a distance matrix ***M*** (for 3 subsequences, ***M*** is a 3×3 matrix) based on the BLAST report for all subsequence alignments. Each element ***M_ij_*** of ***M*** is calculated as follows:

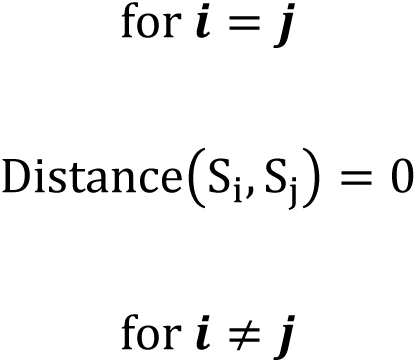

Distance(S_i_, S_j_)

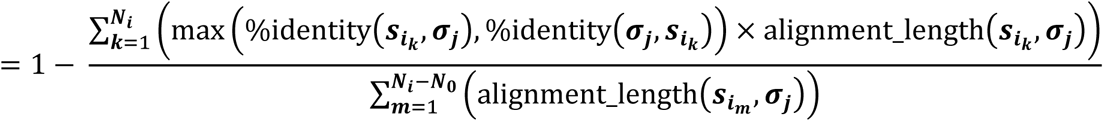

c. The final result is a distance matrix containing the metric described in item b. Various methodologies can then be used to derive a phylogenetic tree based on this matrix (e.g., neighbor joining, weighted least squares, etc.). The distance matrix can be represented as follows:

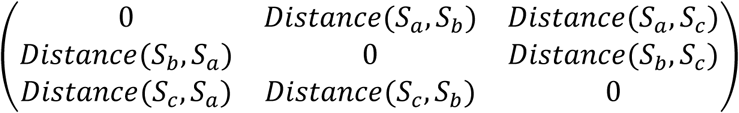

### Dataset of cyanobacterial NRPS/PKS gene clusters

The sequences of toxin-producing gene clusters were obtained from the Nucleotide database of NCBI (queried in February 2018) using the search terms ((NRPS) OR (PKS) OR (gene_cluster) OR (polyketide_synthase) OR (nonribosomal_peptide_synthase)) AND “cyanobacteria”[porgn: txid1117]. Subsequently, we filtered out all sequences shorter than 1,500 bp, and manually removed all entries whose annotation did not refer to NRPS or PKS proteins (contig, scaffold, whole genome, plasmid, etc.), entries containing patent sequences, and entries that did not specify the final product of the gene cluster. To ensure that all relevant entries were identified, we performed additional searches using the identified gene cluster products as keywords (e.g., microcystin, cylindrospermopsin), and applied the same filtering protocol. For each hit, we downloaded a FASTA file containing the coding sequences (CDS), and we labeled each file with its corresponding GenBank accession code. Each file was downloaded and inspected separately. If a file contained also sequences annotated to code for proteins unrelated to the gene cluster product, we removed these particular CDS. If a file contained several regions annotated as different clusters (i.e., resulting in different products), we moved the CDS for each different cluster to a separate file. Finally, we retained only files that contained at least three genes. In these files, we concatenated the remaining sequences by removing non-sequence information (FASTA headers), without changing the order of genes in the original file. The final analysis included only toxins for which at least two biosynthetic gene clusters satisfied the above-listed conditions (19 toxins, 76 biosynthetic gene clusters). Thus, the phylogenetic analysis was performed on a dataset of 76 files, each pertaining to a single biosynthetic cluster and being referred to using a single identifier (e.g., “Microcystin_1”) (Figure 3). Prior to sequence alignment, ambiguous nucleotides were replaced as follows: *n* and *v* were replaced with *a*; *s* and *b* were replaced with *c*; *r*, *m*, and *d* were replaced with *g*; *k*, *w*, *h*, and *u* were replaced with *t*. The dataset (nucleotide sequences in FASTA format) is available as Supplementary Data S1, whereas the phylogenetic tree (in Newick format) is available as Supplementary Data S2.

**Figure 3.**
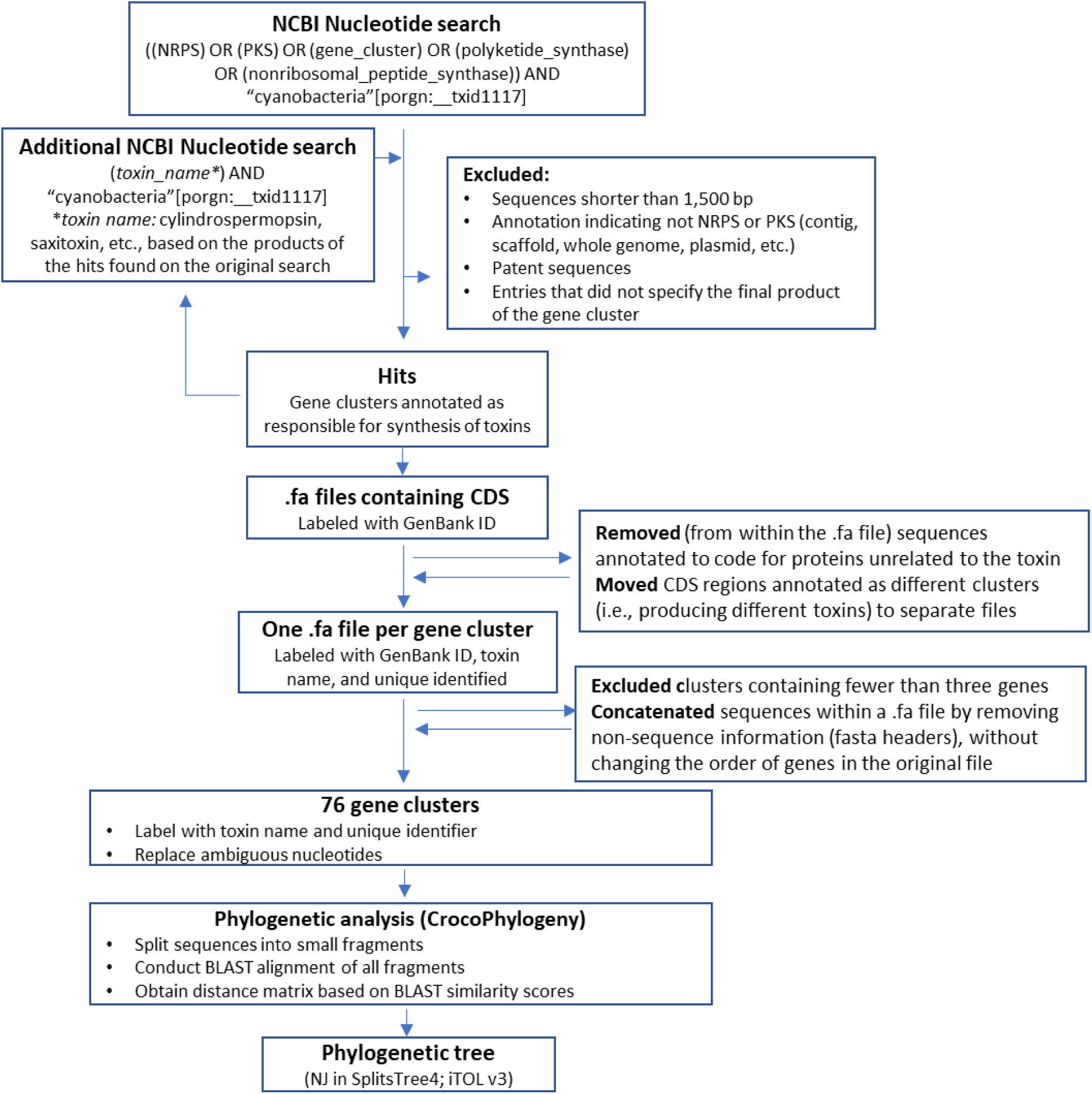
Overview of methods. Full information about the 76 gene clusters included in the analysis is provided in Table S1. The dataset (nucleotide sequences in FASTA format) is available as Supplementary Data S1, whereas the phylogenetic tree (in Newick format) is available as Supplementary Data S2. CDS, coding sequences; PKS, polyketide synthesis; NJ, neighbor joining; NRPS, non-ribosomal peptide synthesis.

### Software and hardware used

The datasets were obtained and processed manually. The phylogenetic analysis was conducted using CrocoPhylogeny version 0.0.18.6.30 with default settings^61^, running NCBI BLAST version 2.6^30^. The phylogenetic tree was built using neighbor joining as implemented in SplitsTree4^62^, with default parameters, starting from the distance matrix. All calculations were performed on a computer with the following specifications: Intel Core i7-5820K CPU (6 cores, 12 threads); 32.0 GB RAM (DDR4); ext4 file system on SSD; Ubuntu 16.4. The phylogenetic tree was visualized using iTOL v3^63^.

## Data availability

All data generated or analyzed during this study are included in this published article and its supplementary information files.

## Acknowledgements

The authors thank Dr. Luísa Hoffmann, Mr. Taraka Ramji Moturu, and Mrs. Michelle Almeida da Paz for helpful comments and useful discussions.

## Author contributions

RJTR, CMI, and JK designed the study. RJTR and CMI obtained the source data and analyzed the results. RJTR wrote the code and ran the calculations. CMI wrote the manuscript. All authors contributed substantially to discussion of the content and reviewed and edited the manuscript before submission.

## Additional information

### Ethics approval and consent to participate

Not applicable, as no human or animal studies were conducted. All data for analysis were obtained from publicly available databases.

### Consent for publication

Not applicable.

## Supplementary information legends

**Supplementary Table S1. Overview of biosynthetic gene clusters used in this study.** Table containing cyanotoxin name, GenBank ID, organism, length, and list of genes in each cluster.

**Supplementary Data S1. Nucleotide sequences of the biosynthetic gene clusters.** The FASTA sequences of all biosynthetic gene clusters are provided in a single file for easy download.

**Supplementary Data S2. Phylogenetic tree of biosynthetic gene clusters.** Phylogenetic tree (Newick format) in which branch lengths are derived from the average pairwise similarity of subsequence alignments.

