## Supplementary Table S1 for "The evolutionary advantage of toxin production among cyanobacteria, the oldest known organisms on Earth"

### Supplementary Table S1: Overview of biosynthetic gene clusters used in this study

| ID, type | GenBank ID | Organism | Length | Genes |
|---|---|---|---|---|
| **Modified cyclic peptides** | | | | |
| Anabaenopeptin_1 | KU665241.1 | Planktothrix agardhii No758 | 23715 | apnA apnB apnC apnD apnE |
| Anabaenopeptin_2 | KU665240.1 | Planktothrix agardhii No365 | 23769 | anaE anaD anaC anaB anaA |
| Anabaenopeptin_3 | KU665237.1 | Planktothrix agardhii NIVA-CYA 126/8 | 23763 | aptA aptB aptC aptD aptE aptF |
| Anabaenopeptin_4 | KU665238.1 | Planktothrix agardhii No66 | 23712 | AptA AptB AptC AptD AptE AptF |
| Anabaenopeptin_5 | AM990463.2 | Planktothrix prolifica NIVA-CYA 98 | 23715 | aptA1 aptA2 aptB aptC aptD aptE aptF |
| Anabaenopeptin_6 | KX788858.1 | Sphaerospermopsis torques-reginae ITEP-024 | 25407 | apnA apnB apnC apnD apnE |
| Anabaenopeptin_7 | MF196967.1 | Nostoc sp. KVJ2 | 25080 | apnA apnB apnC apnD apnE |
| Anabaenopeptin_8 | GU174493.1 | Anabaena sp. 90 | 31710 | apnA apnB apnC apnD apnE |
| Anabaenopeptin_9 | KU665236.1 | Planktothrix rubescens NIVA-CYA 18 | 23715 | apnA apnB apnC apnD apnE |
| Anabaenopeptin_10 | KU665239.1 | Planktothrix rubescens No82 | 23715 | apnA apnB apnC apnD apnE |
| Anabaenopeptin_11 | KU665235.1 | Planktothrix rubescens No108 | 23742 | apnA apnB apnC apnD apnE |
| Anabaenopeptin_12 | EF672686.1 | Planktothrix agardhii NIVA-CYA 126/8 | 23763 | apnA apnB apnC apnD apnE |
| Anacyclamide_1 | FJ461733.2 | Anabaena sp. 90 | 5544 | acyC acyB acyA acyE acyF acyG |
| Anacyclamide_2 | MF140698.1 | Sphaerospermopsis sp. LEGE 00249 | 3414 | acyC acyB acyA acyE acyF |
| Cyanopeptolin_1 | KU665241.1 | Planktothrix agardhii No758 | 20589 | ociC ociB ociD |
| Cyanopeptolin_2 | KU665240.1 | Planktothrix agardhii No365 | 20733 | ociC ociB ociD |
| Cyanopeptolin_3 | KU665237.1 | Planktothrix agardhii NIVA-CYA 126/8 | 27735 | ociC ociB ociA ociD |
| Cyanopeptolin_4 | KU665238.1 | Planktothrix agardhii No66 | 20469 | ociC ociB ociD |
| Cyanopeptolin_5 | AM990463.2 | Planktothrix prolifica NIVA-CYA 98 | 32880 | ociD ociA ociB ociG ociH ociC |
| Cyanopeptolin_6 | DQ837301.1 | Planktothrix agardhii NIVA-CYA 116 | 30939 | ociD ociA ociB ociC |
| Cyanopeptolin_7 | DQ075244.1 | Microcystis sp. NIVA-CYA 172/5 | 30156 | mcnG mcnF mcnA mcnB mcnC mcnD mcnE |
| Microcyclamide_1 | AM774406.1 | Microcystis aeruginosa NIES-298 | 10047 | mcaA mcaB mcaC mcaD mcaE mcaF mcaG |
| Microcyclamide_2 | AM931579.1 | Microcystis aeruginosa PCC 7806 | 10017 | mcaA mcaB mcaC mcaD mcaE mcaF mcaG |
| Microcystin_1 | KC699835.1 | Nostoc sp. 152 (Nostoc sp. PCC 9237/1) | 52683 | mcyH mcyI mcyF mcyE mcyD mcyG mcyA mcyB mcyC mcyL |
| Microcystin_2 | AF183408.1 | Microcystis aeruginosa PCC 7806 | 52950 | mcyJ mcyI mcyH mcyG mcyF mcyE mcyD mcyA mcyB mcyC |
| Microcystin_3 | AY212249.1 | Anabaena sp. 90 | 53076 | mcyH mcyI mcyF mcyE mcyJ mcyD mcyG mcyA mcyB mcyC |
| Microcystin_4 | AJ536156.1 | Anabaena sp. 90 | 53076 | mcyH mcyI mcyF mcyE mcyJ mcyD mcyG mcyA mcyB mcyC |
| Microcystin_5 | AJ441056.1 | Planktothrix agardhii NIVA-CYA 126/8 | 52272 | mcyT mcyD mcyE mcyG mcyH mcyA mcyB mcyC mcyJ |
| Microcystin_6 | AM990462.1 | Planktothrix prolifica NIVA-CYA 98 | 49434 | mcyD mcyE mcyG mcyH mcyA mcyB mcyC |
| Microcystin_7 | AB032549.2 | Microcystis aeruginosa | 34461 | mcyD mcyE mcyF mcyG mcyH mcyI mcyJ |
| Nodularin_1 | AY210783.2 | Nodularia spumigena strain NSOR10 | 46164 | ndaB ndaA ndaC ndaD ndaE ndaF ndaG ndaH ndaI |
| Nodularin_2 | MF668122.1 | Nostoc sp. CENA543 | 46422 | ndaI ndaH ndaG ndaF ndaE ndaD ndaC ndaA ndaB |
| **Modified amino acids** | | | | |
| Mycosporine_1 | KX021865.1 | Scytonema cf. crispum UCFS10 | 7500 | mysA mysB mysC mysE |
| Mycosporine_2 | KX021866.1 | Scytonema cf. crispum UCFS15 | 7500 | mysA mysB mysC mysE |
| **Modified non-cyclic peptides with alkaloid moiety** | | | | |
| Aeruginosin_1 | AM990465.1 | Planktothrix rubescens | 23271 | aerA aerB aerC aerD aerE aerF aerG aerH aerI aerK aerL aerJ |
| Aeruginosin_2 | FJ609416.1 | Microcystis aeruginosa | 22497 | aerA aerJ aerB aerK aerD aerE aerF aerL aerG1 aerG2 aerM aerN |
| **Alkaloids** | | | | |
| Ambiguine_1 | KJ742065.1 | Fischerella ambigua UTEX 1903 | 34635 | ambS1 ambS2 ambC3 ambT5 ambT4 ambT3 ambT2 ambT1 ambC2 ambD3 ambD2 ambP1 ambI3 ambI2 ambI1 ambD1 ambC1 ambR2 ambR1 ambP2 ambD4 ambP3 ambO1 ambO2 ambO3 ambU1 ambU2 ambU3 ambU4 ambO4 ambO5 ambE1 ambE2 ambE3 ambO6 ambO7 |
| Ambiguine_2 | KF664586.1 | Fischerella ambigua UTEX 1903 | 31812 | AmbE3 AmbE2 AmbE1 AmbO5 AmbO4 AmbU4 AmbU3 AmbU2 AmbU1 AmbO3 AmbO2 AmbO1 AmbP3 AmbD4 AmbP2 AmbR1 AmbR2 AmbC1 AmbD1 AmbI1 AmbI2 AmbI3 AmbP1 AmbD2 AmbD3 AmbC2 AmbT1 AmbT2 AmbT3 AmbT4 AmbT5 AmbC3 |
| Ambiguine_3 | KX451322.1 | Fischerella ambigua UTEX 1903 | 31872 | famA1 famA2 famA3 famB1 famB2 famC1 famC2 famC3 famC4 famB3 famB4 famB5 famD1 famE1 famE2 famF1 famF2 famG1 famE3 famH1 famH2 famH3 famD2 famE4 famE5 famI1 famI2 famI3 famI4 famI5 famI6 famG2 |
| AnatoxinA_1 | FJ477836.2 | [Oscillatoria] sp. PCC 6506 (Oscillatoria sp. PCC 9029 = Lyngbya kuetzingii UTEX B 1547) | 23316 | anaI anaJ anaA anaB anaC anaD anaE anaF anaG anaH |
| AnatoxinA_2 | JF803645.1 | Anabaena sp. 37 | 20736 | anaB anaC anaD anaE anaF anaG anaA |
| AnatoxinA_3 | KM245024.1 | Cuspidothrix issatschenkoi RM-6 | 20787 | anaA anaB anaC anaD anaE anaF anaG |
| AnatoxinA_4 | KM245023.1 | Cuspidothrix issatschenkoi LBRI48 | 20787 | anaA anaB anaC anaD anaE anaF anaG |
| AnatoxinA_5 | KM245025.1 | Cuspidothrix issatschenkoi CHABD3 | 19110 | anaA anaB anaC anaD anaE anaF anaG |
| Barbamide_1 | AF516145.1 | Lyngbya majuscula 19L | 25155 | barA barB1 barB2 barC barD barE barF barG barH barI barJ barK |
| Barbamide_2 | HQ696501.1 | Moorea producens 3L | 25155 | BarK BarJ BarI BarH BarG BarF BarE BarD BarC BarB2 BarB1 BarA |
| CuracinA_1 | AY652953.1 | Lyngbya majuscula strain 19L | 63345 | curA curB curC curD curE curF curG curH curI curJ curK curL curM curN |
| CuracinA_2 | HQ696500.1 | Moorea producens 3L | 62442 | CurM CurL CurK CurJ CurI CurH CurG CurF CurE CurD CurC CurB CurA |
| Cylindrospermopsin_1 | GQ385961.1 | Aphanizomenon sp. 10E6 | 36156 | cyrJ cyrH cyrK cyrI cyrG cyrF cyrD cyrC cyrA cyrB cyrE |
| Cylindrospermopsin_2 | KJ139743.1 | Cylindrospermopsis raciborskii CHAB3438 | 36147 | cyrA cyrB cyrE cyrC cyrD cyrF cyrG cyrI cyrK cyrH cyrJ |
| Cylindrospermopsin_3 | FJ418586.4 | [Oscillatoria] sp. PCC 6506 (Oscillatoria sp. PCC 9029 = Lyngbya kuetzingii UTEX B 1547) | 36420 | cyrD cyrC cyrH cyrK cyrA cyrB cyrE cyrJ cyrF cyrG cyrI |
| Cylindrospermopsin_4 | JN873921.1 | Raphidiopsis curvata CHAB1150 | 35358 | cyrA cyrB cyrE cyrC cyrD cyrF cyrG cyrK cyrH cyrJ |
| Cylindrospermopsin_5 | KJ139745.1 | Raphidiopsis curvata HB1 | 35358 | cyrA cyrB cyrE cyrC cyrD cyrF cyrG cyrK cyrH cyrJ |
| Cylindrospermopsin_6 | EU140798.1 | Cylindrospermopsis raciborskii AWT205 | 39081 | cyrD cyrF cyrG cyrI cyrK cyrL cyrH cyrJ cyrA cyrB cyrE cyrC cyrN cyrO |
| Cylindrospermopsin_7 | KJ139742.1 | Cylindrospermopsis raciborskii CHAB358 | 35316 | cyrA cyrB cyrE cyrC cyrD cyrF cyrG cyrK cyrH cyrJ |
| Fischerindole_1 | KY026488.1 | Fischerella sp. TAU | 32946 | filA1 filA2 filA3 filB1 filB2 filC1 filB3 filC2 filC3 filC4 filB4 filB5 filB6 filD1 filE1 filE2 filF1 filF2 filG1 filE3 filH1 filH2 filH3 filD2 filE4 filE5 filI1 filI2 filI3 filI4 filI5 filI6 filG2 |
| Fischerindole_2 | KF819511.1 | Fischerella muscicola UTEX 1829 | 29646 | FidE3 FidE2 FidE1 FidO4 FidO3 FidU4 FidU3 FidU2 FidO2 FidO1 FidU1 FidD4 FidP2 FidR1 FidR2 FidC1 FidD1 FidI1 FidI2 FidI3 FidP1 FidD2 FidD3 FidC2 FidT1 FidT2 FidT3 FidT4 FidT5 FidC3 |
| Fischerindole_3 | KY026487.1 | Fischerella muscicola UTEX 1829 | 30054 | fimA1 fimA2 fimA3 fimB1 fimB2 fimC1 fimC2 fimC3 fimC4 fimJ fimB3 fimC5 fimE1 fimE2 fimF1 fimF2 fimG1 fimE3 fimH1 fimH2 fimH3 fimD fimE4 fimE5 fimI1 fimI2 fimI3 fimI4 fimI5 fimI6 fimG2 |
| NitFischerindole_1 | KY026489.1 | Fischerella sp. SAG 46.79 | 24708 | fisB1 fisB2 fisC fisE1 fisE2 fisM fisG1 fisE3 fisH1 fisH2 fisH3 fisD fisE4 fisE5 fisB3 fisB4 fisI1 fisI2 fisI3 fisI4 fisI5 fisI6 fisG2 |
| NitFischerindole_2 | KY681023.1 | Fischerella sp. SAG 46.79 | 24708 | NifO5 NifO1 NifU1 NifD4 NifP2 NifM NifC1 NifD1 NifI1 NifI2 NifI3 NifP1 NifD2 NifD3 NifO2 NifO3 NifC2 NifT1 NifT2 NifT3 NifT4 NifT5 NifC3 |
| NocuolinA_1 | KP143720.1 | Nostoc sp. CCAP 1453/38 | 36150 | nocA nocB nocC nocD nocE nocF nocG nocH nocI nocJ nocK nocL nocM nocN nocO nocP nocQ nocR nocS nocT |
| NocuolinA_2 | KY594676.1 | Nodularia sp. HBU26 | 33840 | nocA nocB nocE nocF nocG nocH nocI nocJ nocK nocL nocC nocM nocO nocP nocQ nocR nocS nocT |
| Saxitoxin_1 | EU603711.1 | Microseira wollei (Lyngbya) | 27294 | sxtA sxtB sxtC sxtPER sxtN1 sxtM1 sxtG sxtH sxtM2 sxtI sxtACT sxtSUL sxtDIOX sxtM3 sxtX sxtW sxtV sxtU sxtT sxtS sxtR sxtQ sxtP sxtD sxtE |
| Saxitoxin_2 | DQ787200.1 | Cylindrospermopsis raciborskii T3 | 26415 | sxtD sxtC sxtB sxtA sxtE sxtF sxtG sxtH sxtI sxtJ sxtK sxtL sxtM sxtN sxtX sxtW sxtV sxtU sxtT sxtS sxtR sxtQ sxtP sxtO sxtY sxtZ |

| | | | | | | | | | | | | | | | | | | | | | | | | | | | | | | | | | | | | | | | | |
|---|---|---|---|---|---|---|---|---|---|---|---|---|---|---|---|---|---|---|---|---|---|---|---|---|---|---|---|---|---|---|---|---|---|---|---|---|---|---|---|---|
| Saxitoxin_3 | DQ787201.1 | Dolichospermum circinale AWQC131C (Anabaena circinalis AWQC131C) | 19410 | sxtB | sxtA | sxtE | sxtD | sxtP | sxtR | sxtS | sxtT | sxtU | sxtN | sxtG | sxtH | sxtM | sxtI | sxtJ | sxtK | sxtL | sxtO | | | | | | | | | | | | | | | | | | | |
| Saxitoxin_4 | EU603710.1 | Aphanizomenon sp. NH-5 | 21864 | sxtPER | sxtC | sxtB | sxtA | sxtE | sxtW | sxtX | sxtD | sxtP | sxtQ | sxtR | sxtS | sxtT | sxtU | sxtN | sxtG | sxtH | sxtM | sxtI | sxtJ | sxtK | sxtL | | | | | | | | | | | | | | | |
| Saxitoxin_5 | LT549448.1 | Aphanizomenon gracile NIVA-CYA 851 | 21858 | sxtPER | sxtC | sxtB | sxtA | sxtE | sxtW | sxtX | sxtD | sxtP | sxtQ | sxtR | sxtS | sxtT | sxtU | sxtN | sxtG | sxtH | sxtM | sxtI | sxtJ | sxtK | sxtL | | | | | | | | | | | | | | | |
| Saxitoxin_6 | LT549446.1 | Aphanizomenon gracile NIVA-CYA 655 | 21864 | sxtPER | sxtC | sxtB | sxtA | sxtE | sxtW | sxtX | sxtD | sxtP | sxtQ | sxtR | sxtS | sxtT | sxtU | sxtN | sxtG | sxtH | sxtM | sxtI | sxtJ | sxtK | sxtL | | | | | | | | | | | | | | | |
| Saxitoxin_7 | LT549447.1 | Aphanizomenon gracile UAM529 | 21864 | sxtPER | sxtC | sxtB | sxtA | sxtE | sxtW | sxtX | sxtD | sxtP | sxtQ | sxtR | sxtS | sxtT | sxtU | sxtN | sxtG | sxtH | sxtM | sxtI | sxtJ | sxtK | sxtL | | | | | | | | | | | | | | | |
| Saxitoxin_8 | LT549449.1 | Aphanizomenon gracile NIVA-CYA 676 | 21864 | sxtPER | sxtC | sxtB | sxtA | sxtE | sxtW | sxtX | sxtD | sxtP | sxtQ | sxtR | sxtS | sxtT | sxtU | sxtN | sxtG | sxtH | sxtM | sxtI | sxtJ | sxtK | sxtL | | | | | | | | | | | | | | | |
| Welwitindolinone_1 | KJ767018.1 | Westiella intricata UH HT-29-1 | 38592 | welO15 | welM1 | welU8 | welO14 | welO13 | welO12 | welO11 | welU6 | welD4 | welP2 | welR1 | welR2 | welR3 | welC3 | welT5 | welT4 | welT3 | welT2 | welT1 | welC2 | welO18 | welO19 | welD3 | welD2 | welP1 | welI3 | welI2 | welI1 | welD1 | welC1 | welM2 | welH | welM3 | welE4 | welO16 | welO17 | welS1 |
| Welwitindolinone_2 | KJ767017.1 | Hapalosiphon welwitschii UH IC-52-3 | 35655 | welO15 | welM1 | welU8 | welO14 | welO13 | welO12 | welO11 | welU6 | welU7 | welD4 | welP2 | welR1 | welR2 | welR3 | welC3 | welT5 | welT4 | welT3 | welT2 | welT1 | welC2 | welD3 | welD2 | welP1 | welI3 | welI2 | welI1 | welD1 | welC1 | welM2 | welH | welM3 | welE4 | welO16 | welO17 | | |
| Welwitindolinone_3 | KF811479.1 | Hapalosiphon welwitschii UTEX B 1830 | 29856 | WelO5 | WelM | WelU2 | WelO4 | WelO3 | WelO2 | WelO1 | WelU1 | WelD4 | WelP2 | WelR1 | WelR2 | WelR3 | WelC3 | WelT5 | WelT4 | WelT3 | WelT2 | WelT1 | WelC2 | WelD3 | WelD2 | WelP1 | WelI3 | WelI2 | WelI1 | WelS1 | WelS2 | WelD1 | WelC1 | | | | | | | |

**Cyclic lipopeptides**

| | | | | | | | | |
|---|---|---|---|---|---|---|---|---|
| Anabaenolysin_1 | KP761740.1 | Anabaena sp. XPORK15F | 21276 | ablD | ablF | ablC | ablB | ablA |
| Anabaenolysin_2 | KP761741.1 | Anabaena sp. XPORK13A | 20532 | ablE | ablD | ablC | ablB | ablA |
| Anabaenolysin_3 | KP761742.1 | Anabaena sp. XSPORK2A | 20532 | ablE | ablD | ablC | ablB | ablA |
